# Targeting comorbid diseases via network endopharmacology

**DOI:** 10.1101/313809

**Authors:** Juaquim Aguirre-Plans, Janet Piñero, Jörg Menche, Ferran Sanz, Laura I Furlong, Harald H. H. W. Schmidt, Baldo Oliva, Emre Guney

## Abstract

The traditional drug discovery paradigm has shaped around the idea of “one target, one disease”. Recently, it has become clear that not only it is hard to achieve single target specificity but also it is often more desirable to tinker the complex cellular network by targeting multiple proteins, causing a paradigm shift towards polypharmacology (multiple targets, one disease). Given the lack of clear-cut boundaries across disease (endo)phenotypes and genetic heterogeneity across patients, a natural extension to the current polypharmacology paradigm is targeting common biological pathways involved in diseases, giving rise to “endopharmacology” (multiple targets, multiple diseases). In this study, leveraging powerful network medicine tools, we describe a recipe for first, identifying common pathways pertaining to diseases and then, prioritizing drugs that target these pathways towards endopharmacology. We present proximal pathway enrichment analysis (PxEA) that uses the topology information of the network of interactions between disease genes, pathway genes, drug targets and other proteins to rank drugs for their interactome-based proximity to pathways shared across multiple diseases, providing unprecedented drug repurposing opportunities. As a proof of principle, we focus on nine autoimmune disorders and using PxEA, we show that many drugs indicated for these conditions are not necessarily specific to the condition of interest, but rather target the common biological pathways across these diseases. Finally, we provide the high scoring drug repurposing candidates that can target common mechanisms involved in type 2 diabetes and Alzheimer’s disease, two phenotypes that have recently gained attention due to the increased comorbidity among patients.

## 1 Introduction

Following Paul Ehrlich’s more-than-a-century-old proposition on magic bullets (one drug, one target, one disease), the drug discovery pipeline traditionally pursues a handful of leads identified in vitro based on their potential to bind to target(s) known to modulate the disease [1]. The success of the selected lead in the consequent clinical validation process relies on the prediction of a drug’s effect in vivo. Yet, in practice, due to the interactions of the compound and its targets with other proteins and metabolites in the cellular network, the characterization of drug effect has been a daunting task, yielding high pre-clinical attrition rates for novel compounds [2, 3].

The high attrition rates can be attributed to the immense response heterogeneity across patients, likely to stem from polygenic nature of most complex diseases. Consequently, researchers have turned their attention to polypharmacology, where novel therapies aim to alter multiple targets involved in the pathway cross-talk pertinent to the disease pathology, rather than single proteins [4, 5]. This has given rise to network-based approaches that predict effect of individual drugs [6] as well as drug combinations [7], allowing to reposition compounds for novel indications.

Over the past years, reusing existing drugs for conditions different than their intended indications has emerged as a cost effective alternative to the traditional drug discovery, giving rise to various drug repurposing methods that aim to mimic the most likely therapeutic and safety outcome of candidate compounds based on the similarities between compounds and diseases characterized by high-throughput omics data [8, 9, 10]. Most studies so far, however, have focused on repurposing drugs for a single condition of interest, failing to recognize the cellular, genetic and ontological complexity inherent to human diseases [11, 12]. Indeed, pathway cross-talk plays an important role in modulating pathophysiology of diseases [13] and most comorbid diseases are interconnected to each other in the interactome through proteins belonging to similar pathways [14, 15, 16, 17]. Recent evidence suggests that endophenotypes [18] –shared intermediate pathophenotypes– such as inflammasome, thrombosome, and fibrosome play essential roles in the progression of many diseases [19].

Here, we propose a novel drug repurposing approach, **P**ro**x**imal pathway **E**nrichment **A**nalysis (PxEA), to specifically target intertangled biological pathways involved in the common pathology of complex diseases. We first identify pathways proximal to disease genes across various autoimmune disorders and use PxEA to investigate whether the drugs promiscuously used in these disorders target specifically the pathways associated to one disease or the pathways shared across the diseases. We find several examples of anti-inflammatory drugs where the pathways proximal to the drug targets in the interactome correspond to the pathways that are in common between two autoimmune disorders, pointing to the existence of immune system related endophenotypes. We demonstrate that PxEA is a powerful computational strategy for targeting multiple pathologies involving common biological pathways such as Alzheimer’s disease and type 2 diabetes. Based on these findings, we argue that PxEA paves the way for targeting multiple disease endophenotypes simultaneously, a concept which we refer as *endopharmacology*.

## 2 Results

### 2.1 Pathway proximity captures the similarities between autoimmune disorders

Conventionally, functional enrichment analysis relies on the significance of the overlap between a set of genes belonging to a condition of interest and a set of genes involved in known biological processes (pathways). Using known pathway genes, one can identify pathways associated to the disease via a statistical test (e.g., P-value from Fisher’s exact test for the overlap between genes or z-score comparing the observed number of common genes to the number of genes one would have in common if genes were randomly sampled from the data set). We start with the observation that such approach often misses key biological process involved in the disease due to the limited overlap between the disease and pathway genes. To show that this is the case, we focus on nine autoimmune disorders for which we obtain genes associated to the disease in the literature and we calculate P-values based on the overlap of these genes to the pathway genes for each of the 674 pathways in Reactome database (Fisher’s exact test, *P* ≤ 0.05). The choice of autoimmune disorders is motivated by the fact that these diseases tend to share key biological functions involved in immune and inflammatory response, for which the current annotation is fairly accurate and complete. Intriguingly, Table 1 demonstrates that the overlap between the disease and pathway genes yields less than ten pathways that are significantly enriched in five out of nine diseases, underestimating extensively the pathology of these disease.

**Table 1:**
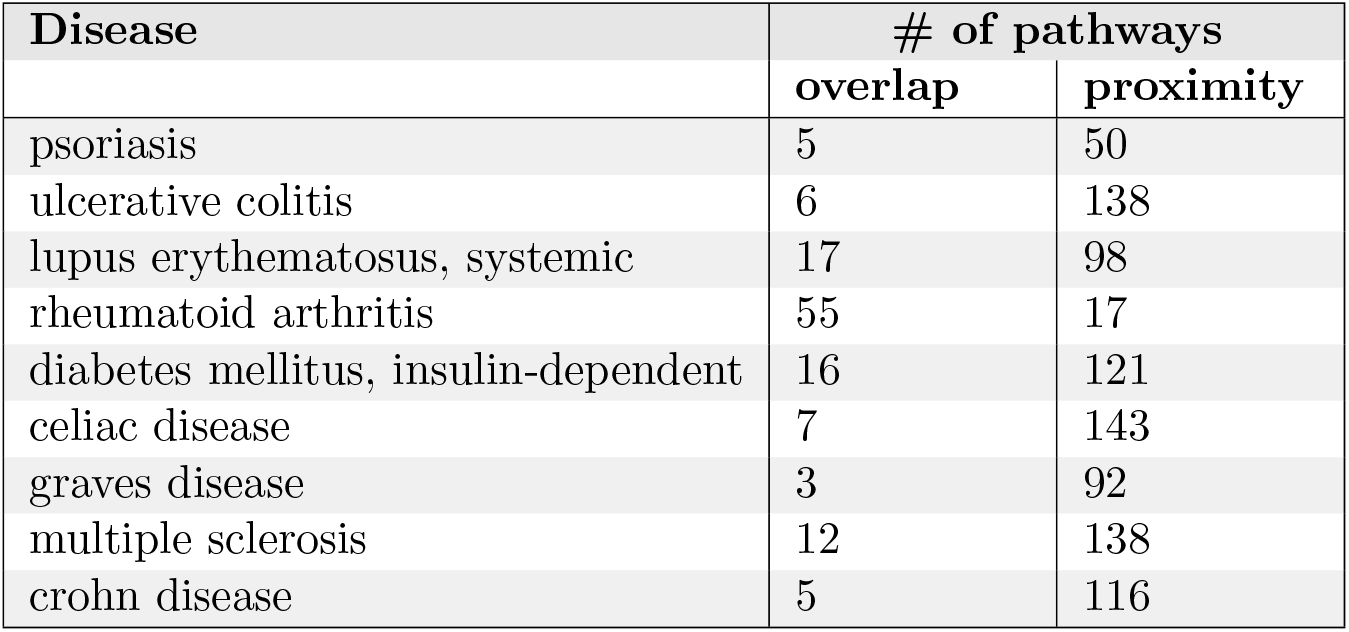
Number of pathways enriched across nine autoimmune disorders based on overlap of pathway and disease genes (*P* ≤ 0.05, assessed by a hypergeometric test) and the proximity of pathway genes to disease genes in the interactome (*z* ≤ −2, see Methods for details).

| Disease | # of pathways |  |
| --- | --- | --- |
|  | overlap | proximity |
| psoriasis | 5 | 50 |
| ulcerative colitis | 6 | 138 |
| lupus erythematosus, systemic | 17 | 98 |
| rheumatoid arthritis | 55 | 17 |
| diabetes mellitus, insulin-dependent | 16 | 121 |
| celiac disease | 7 | 143 |
| graves disease | 3 | 92 |
| multiple sclerosis | 12 | 138 |
| crohn disease | 5 | 116 |

Alternatively, shortest path distance of genes in the interactome can be used to find pathways closer than random expectation to a given set of genes [20, 6], augmenting substantially the number of pathways relevant to the disease pathology. Using network-based proximity [6], we define the *pathway span* of a disease as the set of pathways significantly proximal to the disease (*z* ≤ −2, see Methods). We show that the number of pathways involved in diseases increases substantially when proximity is used (Table 1).

To show the biological relevance of the identified pathways using interactome-based proximity, we check how well these pathways can highlight genetic and phenotypic relationships between nine autoimmune disorders. First, to serve as a background model, we build a disease network for the autoimmune disorders (diseasome) using the genes and symptoms shared between these diseases as well as the comorbidity information extracted from medical claim records (see Methods). The autoimmune diseasome (Figure 1a) is extremely connected covering 33 out of 36 potential links between nine diseases (with average degree < *k* >= 7.3 and clustering coefficient *CC* = 0.93). The three missing links are those in between ulcerative colitis – rheumatoid arthritis, ulcerative colitis – graves disease, and graves disease – type 1 diabetes. On the other hand, several diseases such as celiac disease, crohn disease, systemic lupus erythematosus, and multiple sclerosis are connected to each other with multiple evidence types based on genetic (shared genes) and phenotypic (shared symptoms and comorbidity) similarities, emphasizing the shared pathological components underlying these diseases.

**Figure 1:**
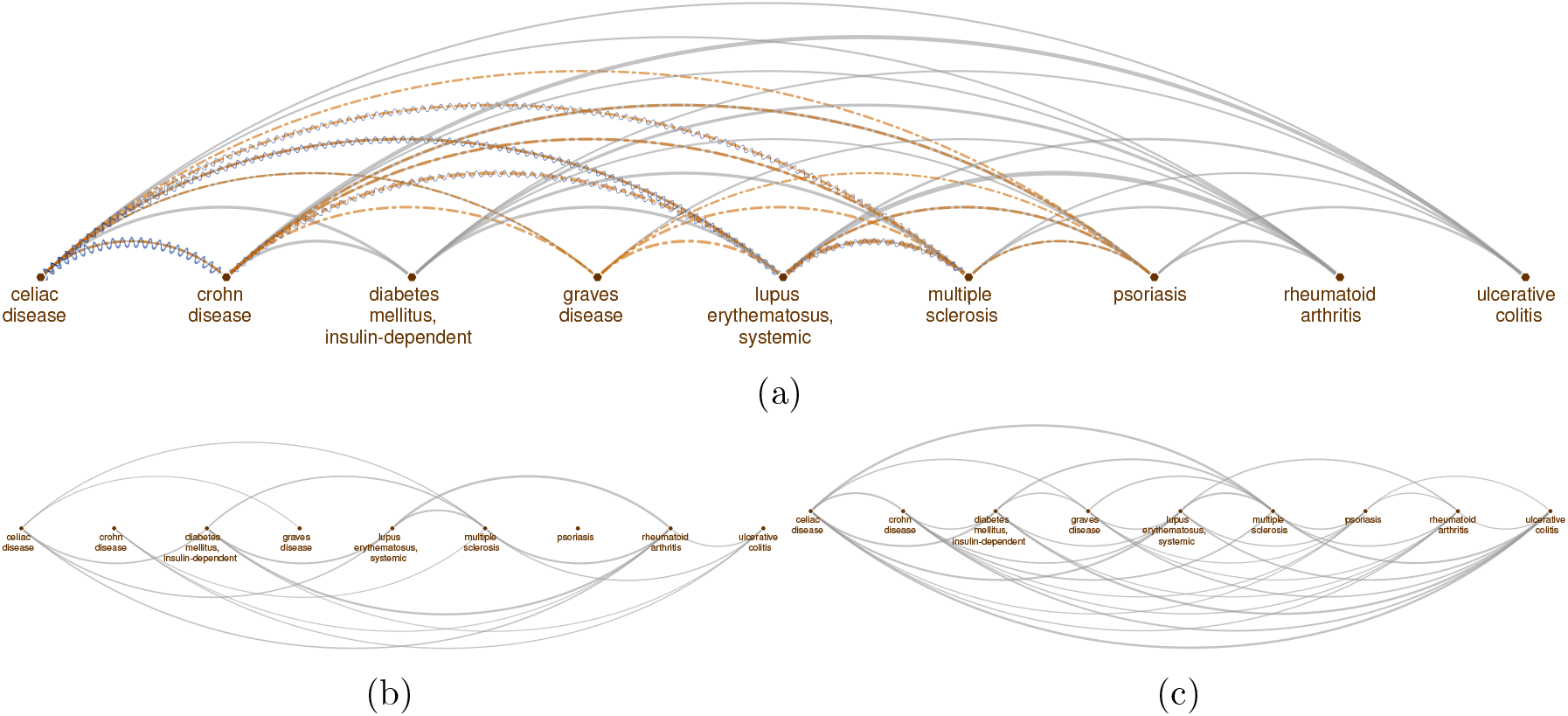
Genetic, phenotypic and functional overlap across autoimmune disorders. Disease relationships (links) based on **(a)** shared genes (gray solid lines), shared symptoms (orange dashed lines) and comorbidity (blue sinusoidal lines), (b) shared pathways (gray solid lines) using common disease and pathway genes, **(c)** shared pathways (gray solid lines) using proximity of pathway genes to diseases genes in the interactome.

We compare the autoimmune diseasome generated using shared genes, common symptoms and comorbidity, to the disease network in which the disease-disease connections are identified using the pathways they share. We identify the pathways enriched in the diseases using both the overlap and proximity approaches mentioned above and check whether the number of common pathways between two diseases is significant (Fisher’s exact test *P* < 0.05). The diseasome based on pathways shared across diseases using overlap between pathway and disease genes is markedly sparser than the original diseasome, containing 17 links (Figure 1b). None of the diseases shares pathways with psoriasis and among the connections supported by multiple evidence in the original diseasome the links between crohn disease and celiac disease as well as crohn disease and systemic lupus erythematosus are missing. On the contrary, the diseasome based on shared pathways using proximity of the pathway and disease genes consists of 34 links, where the only unconnected disease pairs are crohn disease – graves disease and type 1 diabetes – psoriasis, suggesting that it captures the connectedness of the original diseasome better than the overlap-based approach.

We next turn our attention to the shared pathways across diseases identified by both overlap and proximity based approaches and observe that most common pathways involve biological processes relevant to the immune system response through signaling of cytokines (interferon gamma, interleukins such as IL6, IL7) and lymphocytes (ZAP70, PD1, TCR, among others). While overlap based enrichment finds most of these pathways are shared among only 4-5 diseases, proximity based enrichment points to the commonality of these pathways among almost all the diseases. Furthermore, the proximity based enrichment hints involvement of additional interleukin (IL2, IL3, IL5) and lymphocyte (BCR) molecules ubiquitously in autoimmune disorders. These findings suggest that proximity-based pathway enrichment identifies biological processes relevant to the diseases, highlighting the common etiology across autoimmune disorders.

### 2.2 Diseases targeted by the same drugs exhibit functional similarities

Having observed that pathway proximity to diseases in the interactome captures the underlying biological mechanisms across diseases, we seek to investigate the potential implications of the connections between diseases in regards to drug discovery. We hypothesize that a drug indicated for a number of autoimmune diseases would exert its effect by targeting the shared biological pathways across these diseases. To test this, we use 25 drugs that are indicated for two or more of the diseases in Hetionet [21] and split disease pairs into two groups: *(i)* diseases for which a common drug exists and *(ii)* diseases for which no drugs are shared. We then count the number of pathways in common between two diseases for each pair in the two groups using pathway enrichment based on overlap and proximity. We find that the diseases targeted by same drugs tend to involve an elevated number of common pathways compared to the disease pairs that do not have any drug in common (Figure 2). The average number of pathways shared among diseases that are targeted by the same drug is 3.4 and 38 using overlap and proximity based enrichment, respectively, whereas, the remaining disease pairs share 2 and 31 pathways on average using the two enrichment approaches. Though significant only for overlap based approach (*P* = 0.043 and *P* = 0.066 for overlap and proximity, respectively), the difference in the number of pathways is remarkable. We note that due to the relatively small sample size and potentially incomplete drug indication information, we interpret the elevated number of pathways as a trend rather than a general rule across all diseases. Nevertheless, taken together with the high overall pathway-level commonalities observed in the autoimmune disorders mentioned in the previous section, this result suggests that the drugs used for multiple indications are likely to target common pathways involved in these diseases.

**Figure 2:**
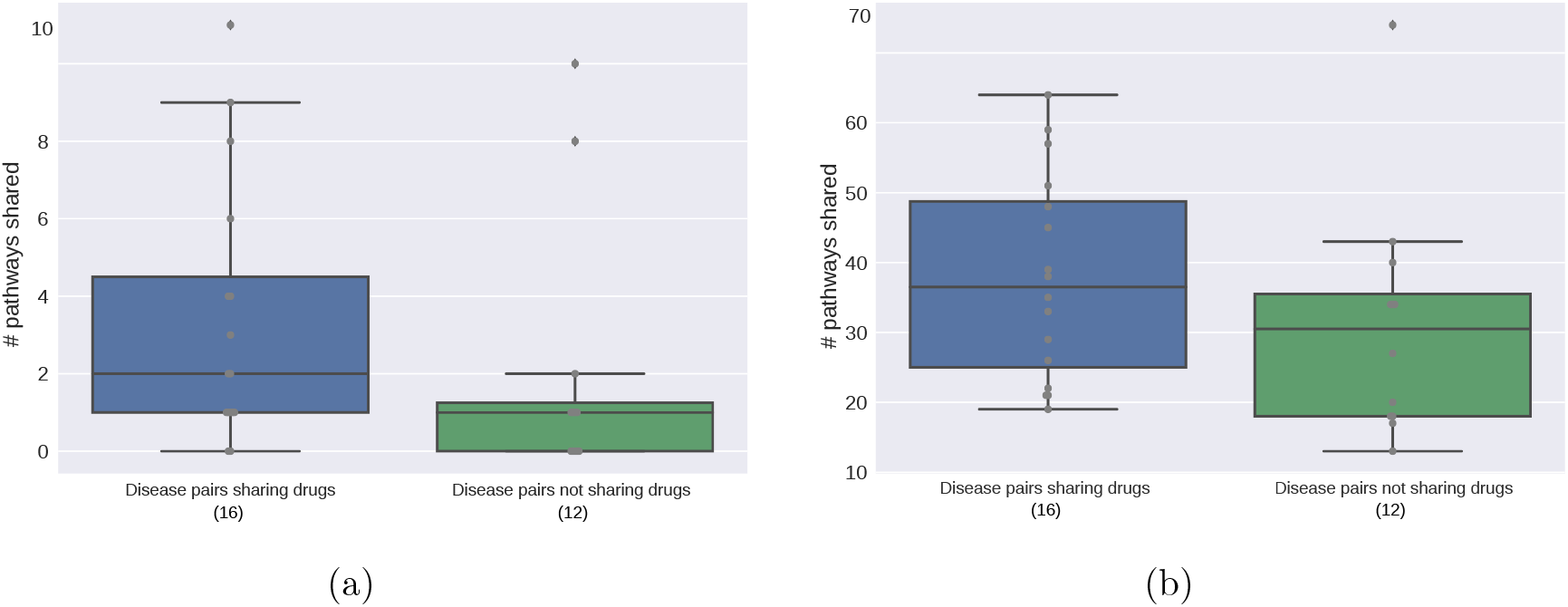
Number of shared pathways across disease pairs that are targeted by the same drug compared to the rest of the pairs. The pathway enrichment is calculated using **(a)** gene overlap and **(b)** proximity of genes in the interactome. The number of disease pairs in each group is given in the parenthesis b below the group label in the x-axis.

### 2.3 Proximal pathway enrichment analysis highlights lack of specificity for drugs used in autoimmune disorders

The observation on the drugs indicated across multiple autoimmune disorders potentially target common pathways triggers us to ask the following question: “Can we leverage pathway-level commonalities between diseases to quantify the impact of a given drug on these diseases?”. To this end, we propose PxEA, a novel method for **P**ro**x**imal pathway **E**nrichment **A**nalysis that scores the likelihood of a set of pathways (e.g., targeted by a drug) to be represented among another set of pathways (e.g., common disease pathways) based on the proximity of the pathway genes in the interactome. As opposed Gene Set Enrichment Analysis (GSEA)[22] which uses gene sets and the ranking of genes based on differential expression, PxEA uses pathway sets and the ranking of pathways based on proximity in the interactome. PxEA scores a drug that involves (i.e. proximal to) multiple pathways in terms of their enrichment among other sets of pathways, such as pathways shared across different diseases. For a given drug and a pair of diseases, we first identify the pathways in the pathway span of both of the diseases, then we rank the pathways with respect to the proximity of the drug targets to the pathway genes and finally we calculate a running sum based statistics corresponding to the enrichment score between the drug and the disease pair (Figure 3, see Methods).

**Figure 3:**
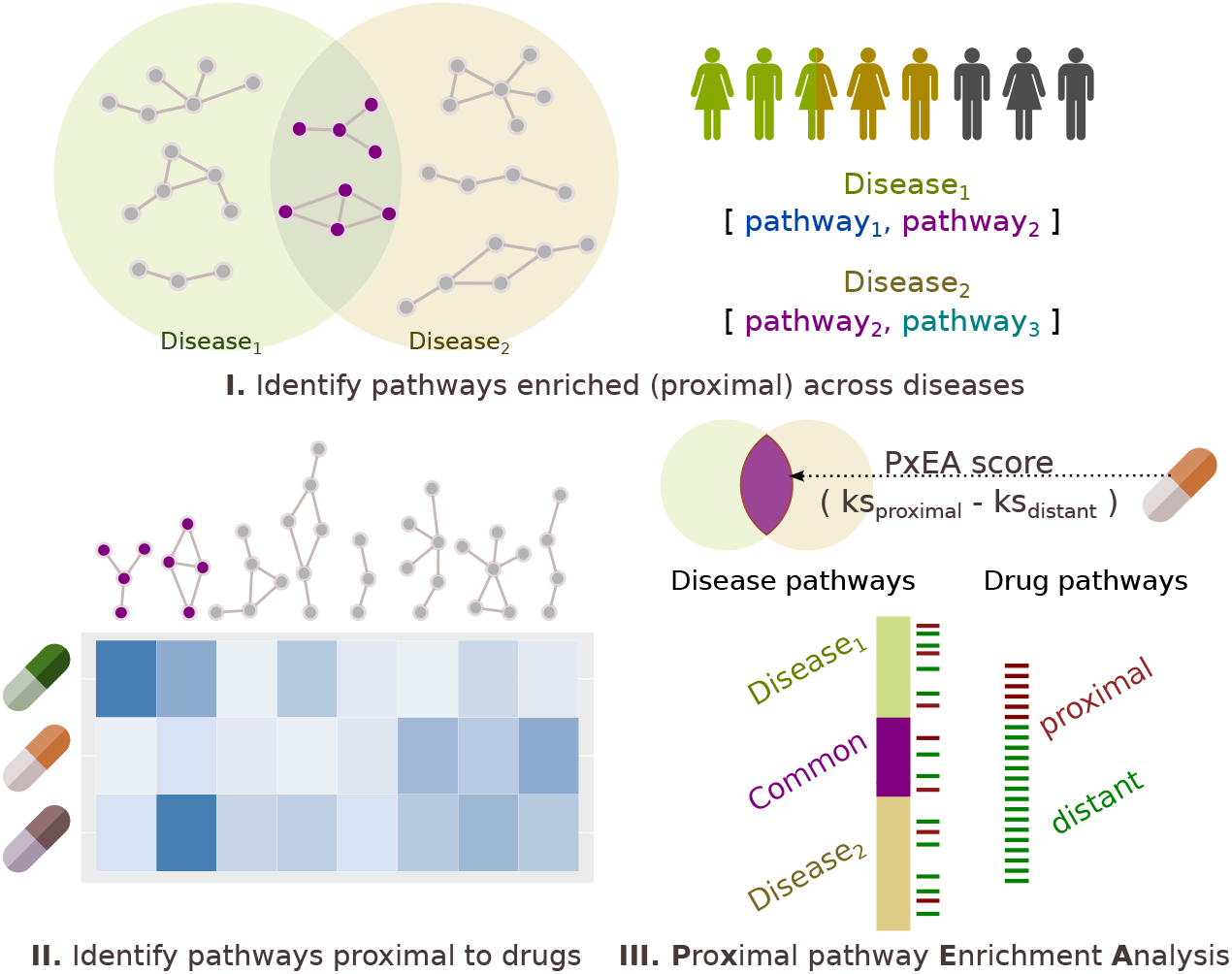
Schematic overview of proximal pathway enrichment analysis (PxEA). PxEA scores a drug with respect to its potential to target the pathways shared between two diseases. For a given drug and two diseases of interest, PxEA first identifies the common pathways between the two disease and then uses the proximity-based ranking of the pathways (i.e. average distance in the interactome to the nearest pathway gene, normalized with respect to a background distribution of expected scores) to assign a score to the drug and the disease pair.

We employ PxEA to score 25 drugs indicated for at least two of the seven autoimmune disorders (there were no common drugs for celiac and graves diseases). For each disease, we first run PxEA using the pathways proximal to the disease and pathways proximal to the drugs that are used for that disease. We then run PxEA for each disease pair, using the pathways proximal to both of the diseases in the pair and the drugs commonly used for the two diseases. We notice that several drugs indicated for multiple conditions score higher using common pathways between two diseases than using pathways of the disease it is indicated for (Figure 4). This is not surprising considering that many of the drugs used for autoimmune disorders target common immune and inflammatory processes. For instance, sildenafil a drug used for the treatment of erectile dysfunction and to relieve symptoms of pulmonary arterial hypertension is reported by Hetionet to show palliative effect on type 1 diabetes and multiple sclerosis. Indeed, sildenafil instead of being specific to any of these two conditions, it targets some of the 57 pathways in common between type 1 diabetes and multiple sclerosis including but not limited to pathways mentioned in Table 2, such as “IL-3, 5 and GM CSF signaling” (*z* = −1.6), “regulation of signaling by CBL” (*z* = −1.1), “regulation of KIT signaling” (*z* = −1.0), “IL receptor SHC signaling” (*z* = 1−1.0), and “growth hormone receptor signaling” (*z* = −1.0).

**Figure 4:**
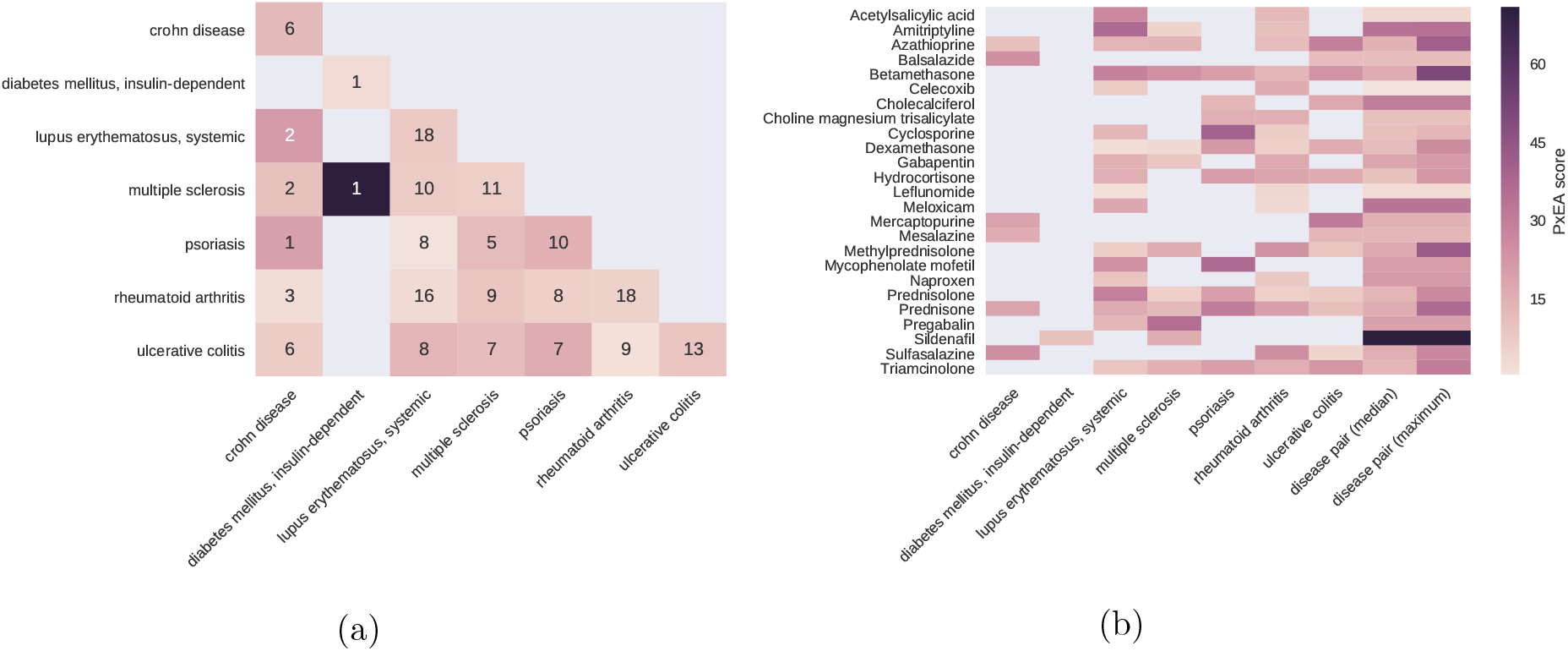
PxEA scores of drugs used in autoimmune disorders. (a) Disease-disease heatmap, in which for each disease pair, the common pathways proximal to the two diseases are used to run PxEA. Note that the diagonal contains the PxEA scores obtained when the proximal pathways for only that disease are used. The hue of the color scales with the PxEA score. (b) Drug-disease heatmap, in which the PxEA is run using the pathways proximal to the pathways of the disease in the column for the drugs in the rows (25 drugs that are used at least in two diseases). The last two columns show the median and maximum values of the PxEA scores obtained for the drug among all disease pairs the drug is indicated for.

**Table 2:** Pathways shared by at least two autoimmune disorders based on gene overlap (hypergeometric *P* ≤ 0.05), pathways shared by at least eight two autoimmune disorders based on proximity of genes and the number of diseases they appear commonly based on both overlap and proximity approaches.

| Pathway | # of shared diseases |  |
| --- | --- | --- |
|  | overlap | proximity |
| interferon gamma signaling | 5 | 8 |
| costimulation by the CD28 family | 5 | 7 |
| cytokine signaling in immune system | 5 | 7 |
| translocation of ZAP-70 to immunological synapse | 5 | 6 |
| phosphorylation of CD3 and TCR zeta chains | 5 | 6 |
| PD1 signaling | 5 | 4 |
| IL-6 signaling | 4 | 8 |
| generation of second messenger molecules | 4 | 6 |
| TCR signaling | 4 | 6 |
| signaling by ILs | 3 | 9 |
| immune system | 3 | 7 |
| downstream TCR signaling | 3 | 7 |
| interferon signaling | 3 | 7 |
| adaptive immune system | 3 | 3 |
| regulation of KIT signaling | 2 | 7 |
| IL-7 signaling | 2 | 6 |
| CTLA4 inhibitory signaling | 2 | 5 |
| chemokine receptors bind chemokines | 2 | 3 |
| extrinsic pathway for apoptosis | 2 | 3 |
| MHC class II antigen presentation | 2 | 2 |
| IL receptor SHC signaling | - | 9 |
| IL-3, 5 and GM CSF signaling | - | 9 |
| signaling by the B cell receptor BCR | - | 8 |
| regulation of IFNG signaling | - | 8 |
| growth hormone receptor signaling | - | 8 |
| IL-2 signaling | - | 8 |
| regulation of signaling by CBL | - | 8 |

Similarly, prednisone, a synthetic anti-inflammatory glucocorticoid agent that is indicated for six of the autoimmune disorders, is assigned a higher PxEA score using pathways shared by crohn disease and systemic lupus erythematosus compared to pathways involved only in ulcerative colitis or in multiple sclerosis, suggesting that this drug does not specifically target any of the autoimmune disorders but rather acts on the common pathology of these diseases. Another example, where we observe a similar trend is meloxicam. While meloxicam is originally indicated for rheumatoid arthritis and systemic lupus erythematosus, the higher PxEA score when common pathways are used suggests that it targets underlying inflammatory processes.

### 2.4 Targeting the common pathology of type 2 diabetes and Alzheimer’s disease

Type 2 diabetes (T2D) and Alzheimer’s disease (AD), two diseases highly prevalent to an ageing society, are known to exhibit increased comorbidity [23, 24]. Recently, repurposing anti-diabetic agents to prevent insulin resistance in AD has gained substantial attention due to the therapeutic potential it offers [25]. Indeed, the pathway spans of T2D and AD covers 170 and 82 Reactome pathways, respectively and 35 pathways are shared among the pathway spans, suggesting a highly significant link between these two diseases in terms of shared pathways (Fisher’s exact test, *P* = 1.3 *×* 10^−45^).

We use PxEA to score 1,466 drugs from DrugBank using the pathways involved in the common pathways of T2D-AD. When we look at the drugs ranked on the top (Table 3), we spot orlistat, a drug indicated for obesity and T2D in Hetionet. Interestingly, existing studies also suggest a role for this drug in the treatment of AD ([26]). Orlistat targets extracellular communication (Ras-Raf-MEK-ERK, NOTCH, and GM-CSF/IL-3/IL-5 signaling) and lipid metabolism pathways (Figure 5). Several of the proteins in these pathways such as APOA1, PSEN2, PNLIP, LPL, and IGHG1 are either themselves Orlistat’s targets or in the close vicinity of the targets that are important for the pathology of the T2D and AD. The second and third top scoring drugs are chenodeoxycholic and obeticholic acid, biliar acids that are in clinical trials for T2D (NCT01666223) and are argued to modulate cognitive changes in AD [27]. We calculate the significance of the PxEA scores by permuting the ranking of the pathways. We find that the adjusted P-values (corrected for multiple hypothesis testing using Benjamini-Hochberg procedure) for the top 10 drugs are all below 0.001, the minimum possible value (due to the 10,000 permutations used in the calculation).

**Table 3:** Top 10 drug repurposing opportunities to target common type 2 diabetes and Alzheimer’s disease pathology.

| Drug | Hetionet indication | PxEA score | Adjusted P-value |
| --- | --- | --- | --- |
| orlistat | obesity, type 2 diabetes | 94.07 | < 0.001 |
| chenodeoxycholic acid | primary biliary cirrhosis | 74.06 | < 0.001 |
| obeticholic acid | - | 74.06 | < 0.001 |
| practolol | - | 70.55 | < 0.001 |
| esmolol | hypertension | 70.55 | < 0.001 |
| clenbuterol | - | 70.44 | < 0.001 |
| erythrityl tetranitrate | - | 70.32 | < 0.001 |
| bupranolol | - | 68.97 | < 0.001 |
| arbutamine | - | 68.97 | < 0.001 |
| fenoterol | - | 68.97 | < 0.001 |

**Figure 5:**
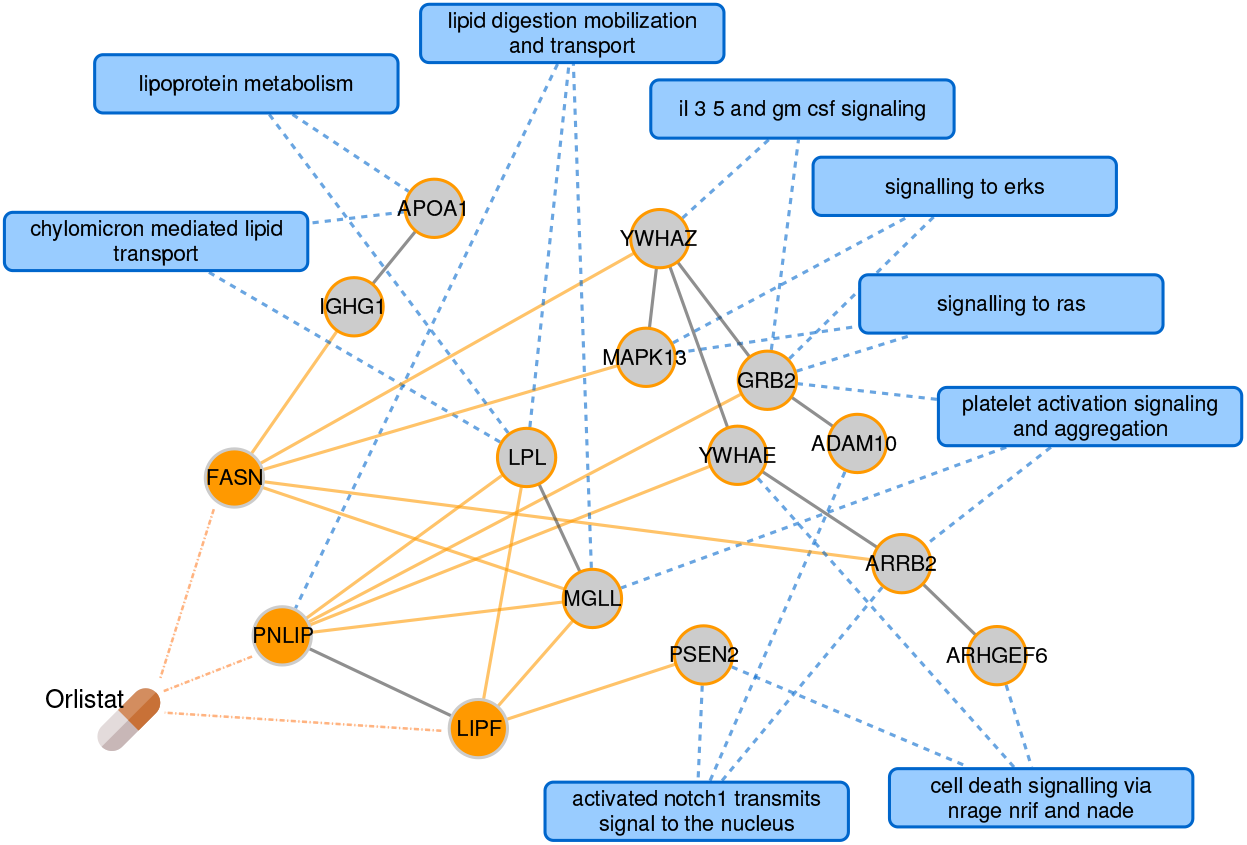
Orlistat from PxEA perspective. The subnetwork shows how the targets of Orlistat are connected to the nearest pathway protein for the pathways shared between T2D and AD. For clarity, only the pathways that are proximal to the drug are shown. Blue rectangles represent pathways, circles represent drug targets (orange) or proteins on the shortest path to the nearest pathway gene (gray). Blue dashed lines denote pathway membership, solid lines are protein interactions. The interactions between the drug and its targets are shown in dashed orange lines and the interactions between the drug targets and their neighbors are highlighted with solid orange lines.

## 3 Discussion

The past decades have witnessed a substantial increase in human life expectancy owing to major breakthroughs in translational medicine. Yet, the increase on average age and changes in life style, have given rise to a spectra of problems challenging human health like cancer, Alzheimer’s disease and diabetes. These diseases do not only limit the life expectancy but also induce a high burden on public healthcare costs. In the US alone, more than 20 and 5 million people have been effected by type 2 diabetes and Alzheimer’s disease, respectively, ranking these diseases among most prevalent health problems [23].

Mainly characterized by hyperglycemia due to resistance to insulin, the disease mechanism of T2D involves a combination of multiple genetic and dietary factors. On the other hand, AD is relatively less understood and several hypothesis have been proposed for its cause: reduced synthesis of neurotransmitter acetylcholine, accumulation of amyloid beta plaques and/or tau protein abnormalities, giving rise to neurofibrillary tangles. Accordingly, most available treatments in AD are palliative (treating symptoms rather than the cause). Given the comorbidity between T2D and AD several studies have recently suggested repurposing diabetes drugs for AD [25]. However, to our knowledge, currently there is no systematic method that can pinpoint drugs that could be useful to target common disease pathology such as the one between T2D and AD.

In this study, we first show that diseases that share drugs also tend to share biological pathways and hypothesize that these pathways can be targeted to exploit novel drug repurposing opportunities. We introduce PxEA, a method based on *(i)* pathways that are proximal to diseases and *(ii)* the ranking of the pathways targeted by a drug using the topology information encoded in the human interactome. We show that PxEA picks up whether drugs target specifically the pathways associated to a disease or common pathways shared across various conditions. We use PxEA to rank drugs for their therapeutic potential in targeting the common disease pathology between T2D and AD. Despite the limitations of PxEA, such as the incompleteness in the drug target, disease and pathway genes, we believe PxEA is the first step towards achieving endopharmacology, that is, targeting common pathways across endophenotypes.

## 4 Methods

### 4.1 Protein interaction data and interactome-based proximity

To define a global map of interactions between human proteins, we obtained the physical protein interaction data from a previous study that integrated various publicly available resources [14]. We downloaded the supplementary data accompanying the article to generate the human protein interaction network (interactome) containing data from MINT[28], BioGRID[29], HPRD[30], KEGG[31], BIGG[32], CORUM[33], PhosphoSitePlus[34]. We used the largest connected component of the interactome in our analyses, which covered 141,150 interactions between 13,329 proteins (represented by ENTREZ gene ids).

Network-based proximity is a graph theoretic approach that incorporates the interactions of a set of genes (i.e. disease genes or drug targets) with other proteins in the human interactome and contextual information as to where the genes involved in pathways reside with respect to the original set of genes [6]. To quantify interactome-based proximity between two gene sets (such as drug targets, pathway genes or disease genes), we used the average shortest path length from the first set to the nearest protein in the second set following the definition in the original study [6]. Accordingly, the proximity from nodes *S* to nodes *T* in a network *G*(*V, E*), is defined as

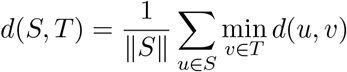

where *d*(*u, v*) is the shortest path length between nodes *u* and *v* in *G*. We then calculated a z-score based on the distribution of the average shortest path lengths across random gene sets *S_random_* and *T_random_* (*d_random_*(*S, T*) = *d*(*S_random_, T_random_*)) as follows:

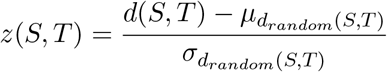

where *μ*_*d_random_*(*S,T*)_ and *σ*_*d_random_*(*T,S*)_ are the mean and the standard deviation of the *d_random_*(*S, T*), respectively obtained using 1,000 realizations of random sampling of gene sets that match the original sets in size and degree. We refer to the pathways that are significantly proximal (*z* ≤ −2) to a disease as the *pathway span* of the disease throughout text.

Note that, instead of average shortest path distances, one can also use random-walk based distances to calculate proximity between gene sets [20]. However, random walks in the networks are inherently biased towards high-degree nodes [35, 36] and require additional statistical adjustment [36, 20]. Sampling based on size and degree matched gene sets has been shown to be robust against data-incompleteness in the interactome and in the known pathway annotations [36, 6]

### 4.2 Disease-gene, drug and pathway information

We compiled genes associated to nine autoimmune disorders listed in Table 4 using disease-gene annotations from DisGeNET [37]. We downloaded curated disease-gene associations from DisGeNET that contained infromation from UniProt [38], ClinVar [39], Orphanet [40], GWAS Catalog [41] and CTD [42]. To ensure that the disease-gene associations were of high confidence, we kept only the associations that were also provided in a previous large-scale analysis of human diseases [14].

**Table 4:**
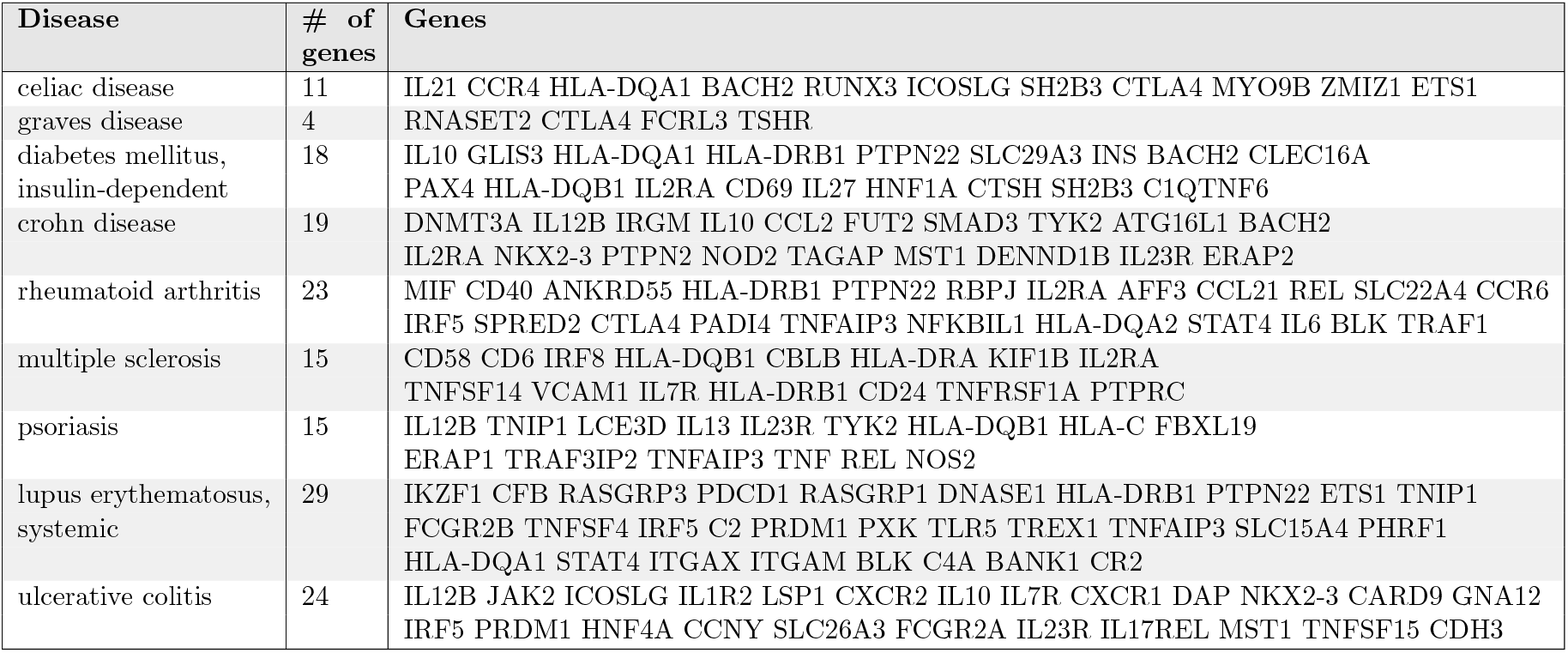
Disease-gene associations for the nine autoimmune disorders used in this study.

We retrieved drug target information from DrugBank for 1,489 drugs in the version 5.0.6 of the database [43], 1,466 of which had at least a target in the interactome. Uniprot ids from DrugBank were mapped to ENTREZ gene ids using Uniprot id mapping file (retrieved on October 2017). We used drug indication information from Hetionet (compound treats or palliates disease edges) that compiled data from publicly available resources [21]. We focused on 78 drugs that were indicated for nine autoimmune disorders above. We created a subset of drugs used for two or more of the autoimmune disorders, yielding 25 drugs across seven conditions (there were no indications for celiac disease, and the two drugs used for graves disease were not used in any other disease).

The ENTREZ gene ids of the proteins involved in biological pathways were taken from the version 5.0 of MSigDB curated gene sets [44]. In our analysis, we used 674 Reactome pathways and the genes associated to these pathways [45].

### 4.3 Genetic, phenotypic and functional relationships across diseases

To identify relationships across disease pairs (autoimmune diseasome), we used the similarities between diseases in terms of genes and symptoms they share. We assessed the significance of the overlap between genes (or symptoms) associated to two diseases using Fisher’s exact test. An alpha value of 0.05 was set to deem the connections significant (*P* ≤ 0.05). The disease symptom information was taken from a previous study based on text mining of PubMed abstracts (only associations with TF-IDF score higher than 3.5 are considered) [46]. We also used the relative risk calculated based on medical insurance claims [47], where we mapped the ICD9 codes to MeSH identifiers using the annotations provided by Disease Ontology [48]. We considered the disease pairs with relative risk higher than 1 as potential commorbidity link.

To identify pathways enriched in diseases, we used the significance *(i)* of the overlap between the pathway and disease genes assessed by a hypergeometic test (i.e. one-sided Fisher’s exact test) and *(ii)* of the proximity between pathway and disease genes in the interactome. We considered the pathways that had *P ≤* 0.05 and *z* ≤ −2, respectively as the pathways that were enriched in a given disease using the two approaches. The pathway information was taken from Reactome and the proximity was calculated as explained above.

### 4.4 PxEA: Proximal pathway enrichment analysis

Toward the goal of pathway-level characterization of the common pathology of diseases and to evaluate the therapeutic potential of drugs based on their impact on the common pathways, we developed **P**roximal pathway **E**nrichment **A**nalysis (PxEA), a novel method that scores drugs based on the proximity of drug targets to pathway genes in the interactome. PxEA uses a GSEA-like running sum score [22], where the pathways are ranked with respect to the proximity of drug targets to the pathways and each pathway is checked whether it appears among the pathways of interest (e.g., common pathways between two diseases). Given *D*, the pathways ranked with respect their proximity to drug targets, *p_i_*, the pathway in consideration within *D* and *C*, the set of pathways of interest, the running score is defined as follows [49]:

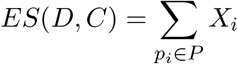

where,

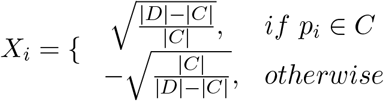

To calculate P-values for the case study, we repeat the procedure above 10,000 times, shuffling randomly *D* to calculate the expected enrichment score *ES*(*D^random^, C*). We then calculate the P-value for the enrichment using

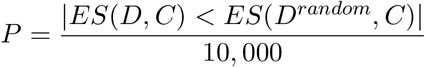

The P-values were corrected for multiple hypothesis testing using Benjamini-Hochberg procedure [50].

### 4.5 Implementation details and code availability

We used toolbox and pxea Python packages for running PxEA, available at http://github.com/emreg00. The proximity was calculated using networkx package [51] that implements Dijkstra’s shortest path algorithm. The statistical tests were conducted in R (http://www.R-project.org) and Python (http://www.python.org). The network visualizations were generated using Cytoscape [52] and the plots were drawn using Seaborn python package [53].

## Acknowledgments

EG is supported by EU-cofunded Beatriu de Pinós incoming fellowship from the Agency for Management of University and Research Grants (AGAUR) of Government of Catalunya and received funding from the Innovative Medicines Initiative 2 Joint Undertaking under grant agreement No 116030. This Joint Undertaking receives support from the European Union’s Horizon 2020 research and innovation programme and EFPIA. L.I.F received support from ISCIII-FEDER (CPII16/00026). The authors also received support from EU H2020 Programme 2014-2020 under grant agreement no. 676559 (Elixir-Excelerate). The Research Programme on Biomedical Informatics (GRIB) is a member of the Spanish National Bioinformatics Institute (INB), PRB2-ISCIII and is supported by grant PT13/0001/0023, of the PE I+D+i 2013-2016, funded by ISCIII and FEDER. The DCEXS is a “Unidad de Excelencia María de Maeztu”, funded by the MINECO (ref: MDM-2014-0370).

